# Real-time experimental control using network-based parallel processing

**DOI:** 10.1101/392654

**Authors:** Byounghoon Kim, Shobha Kenchappa, Adhira Sunkara, Ting-Yu Chang, Lowell Thompson, Raymond Doudlah, Ari Rosenberg

**Author notes:** Correspondence: Ari Rosenberg Department of Neuroscience School of Medicine and Public Health University of Wisconsin – Madison 1111 Highland Ave. WIMR-II, Office 5505 Madison, WI. 53705.

## Abstract

Modern neuroscience research often requires the coordination of multiple processes such as stimulus generation, real-time experimental control, as well as behavioral and neural measurements. The technical demands required to simultaneously manage these processes with high temporal fidelity limits the number of labs capable of performing such work. Here we present an open-source network-based parallel processing framework that eliminates these barriers. The Real-Time Experimental Control with Graphical User Interface (REC-GUI) framework offers multiple advantages: (*i*) a modular design agnostic to coding language(s) and operating system(s) that maximizes experimental flexibility and minimizes researcher effort, (*ii*) simple interfacing to connect measurement and recording devices, (*iii*) high temporal fidelity by dividing task demands across CPUs, and (*iv*) real-time control using a fully customizable and intuitive GUI. Testing results demonstrate that the REC-GUI framework facilitates technically demanding, behavior-contingent neuroscience research. Sample code and hardware configurations are downloadable, and future developments will be regularly released.

## Introduction

Many areas of cutting-edge neuroscience research require real-time experimental control contingent on behavioral and neuronal events, rendering and presentation of complex stimuli, and high-density measurements of neuronal activity. These processes must operate in parallel, and with high temporal resolution. The number of labs that can perform such research is limited by the high technical demands required to set up and maintain an appropriate experimental control system. In particular, a control system must balance the need to precisely coordinate different processes and the flexibility to implement new experimental designs with minimal effort. Systems favoring system precision over usability can hinder productivity because there is a large overhead to learning esoteric or low-level coding languages, and extensive coding demands slow the development of new paradigms. In contrast, systems favoring usability over precision can limit the complexity of supportable paradigms and the ability to perform experiments with high real-time computational demands. Here we present the Real-Time Experimental Control with Graphical User Interface (REC-GUI) framework, which overcomes technical challenges limiting previous solutions by using network-based parallel processing to provide both system precision and usability.

The REC-GUI framework segregates tasks into major groups such as experimental control and monitoring, and stimulus rendering and presentation. Each major group is executed on a different CPU, with communications between CPUs achieved using internet protocols. An additional CPU supports data acquisition and precise temporal alignment of multiple experimental processes. The REC-GUI framework aims to overcome technical challenges that hinder productivity and consume lab resources by providing a solution that works out of the box and reduces the time and effort required to implement experimental paradigms. We have therefore developed a version of the REC-GUI framework in which all processes are implemented with high-level programming environments: experimental control uses a GUI coded in Python, and stimulus rendering and presentation is performed with Psychtoolbox 3 in MATLAB (Brainard, 1997; Pelli, 1997; Kleiner et al., 2007). Psychtoolbox is widely used for its stimulus generation functions that eliminate the need for extensive knowledge of low-level coding languages. The framework is also inherently modular, so system components (e.g., MATLAB for stimulus presentation) can be easily modified or substituted to meet changing research needs. Because the REC-GUI framework achieves precise experimental control with high-level programming environments, it has a low barrier to conducting cutting-edge neuroscience experiments compared to other systems and does not require professional programmers or low-level coding languages. Our testing results confirm that the REC-GUI framework facilitates technically demanding experiments aimed at relating neuronal activity to perception and behavior.

Sample code and hardware configurations providing templates for adaptation and customization are available for download here: https://rosenberg.neuro.wisc.edu/. Developments will be posted with notifications sent through a mailing list. The REC-GUI framework will free time to focus on experimental questions and design, help improve scientific reproducibility by increasing research transparency, and enable scientific advances by reducing technical barriers associated with complex neuroscience studies.

## Materials and methods

### Serial processing, multithreading, and network-based parallel processing alternatives for experimental control

Several approaches can be used to implement experimental control. Here we benchmark different alternatives and identify an option that can satisfy the joint needs of high temporal precision, experimental flexibility, and minimizing coding demands. To illustrate differences between approaches, consider the task of mapping the visual receptive field of a neuron in an awake behaving animal. To perform this task, the animal must hold its gaze on a fixation target presented on a screen. While the animal maintains fixation, the experimenter must control the movement of a visual stimulus such as a bar on the screen. The success of the mapping depends upon the animal maintaining accurate and precise gaze on the target. As such, if the animal breaks fixation, the visual stimulus must disappear until the animal reacquires fixation. The code required to implement this simple task includes three major processes that interface multiple hardware and software components: (*i*) eye position tracker with monitoring routines, (*ii*) visual display for stimulus presentation with functions to account for fixation status, and (iii) interactive experimental control for moving the stimulus, changing stimulus parameters, etc. in real time.

A *serial processing framework* executes these processes serially within a while-loop (***Figure 1A***). As a consequence of serial processing, every iteration of the loop has a pause in eye position monitoring while the stimulus-related and user-related processes finish. This delay can be problematic, especially if the stimulus rendering demands are high. For example, if rendering and presenting the stimulus takes longer than one cycle of the eye position monitoring process, cycles of eye position data will be lost, and the accuracy of the gaze-contingent control compromised. Using this problem as an illustrative example of the challenges that arise for an experimental control system, we next consider how the problem might be solved using a multithreading framework.

**Figure 1.**
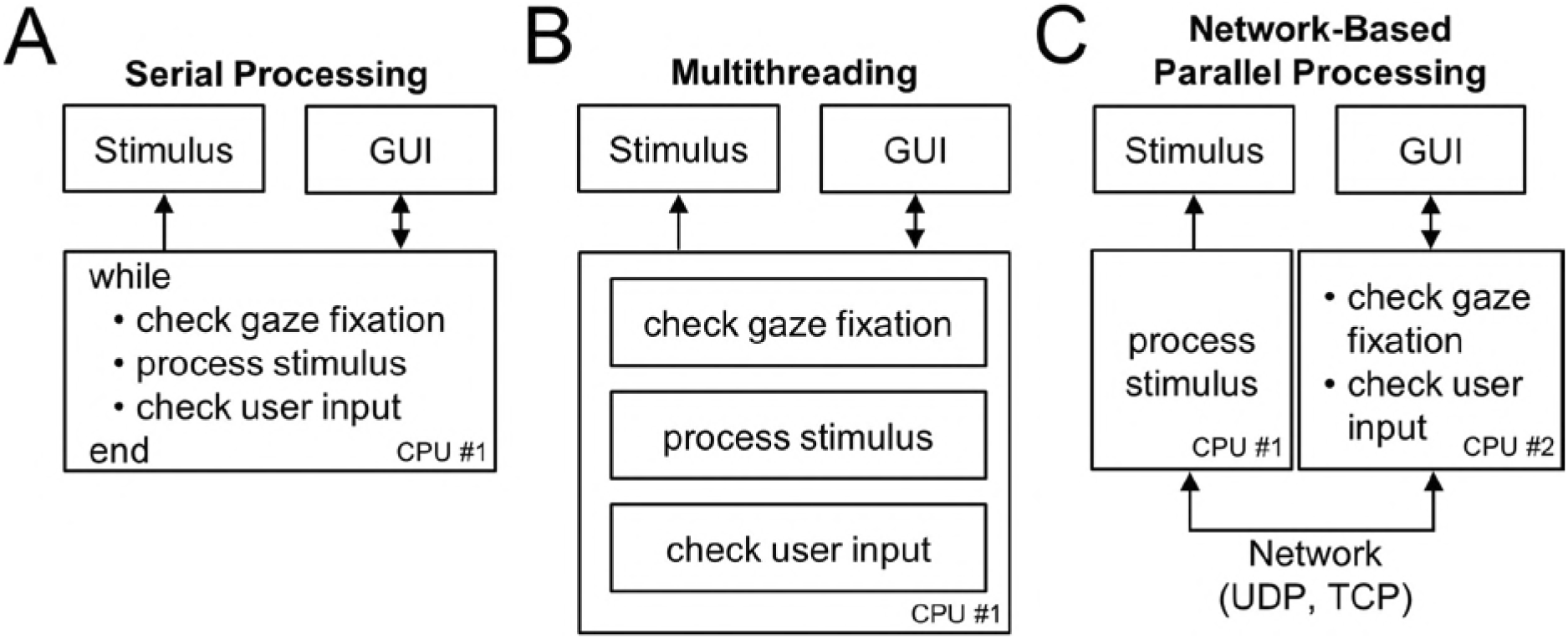
Alternative experimental control frameworks. (A) Serial processing: All processes are executed serially in a while-loop. (B) Multithreading: A single CPU executes multiple processes in parallel on different threads. (C) Network-based parallel processing: Multiple processes are executed in parallel on different CPUs coordinated over a network. Individual processes shown in CPU #2 can be implemented serially or using multithreading. Arrows specify the direction of information flow.

A multithreading framework provides a solution to this problem by allowing the CPU to execute multiple processes concurrently (***Figure 1B***). Specifically, by separating the eye position monitoring, stimulus-related, and user-related processes onto independent threads, a CPU can execute the processes in parallel such that eye position monitoring can proceed without having to wait for the stimulus- and user-related processes to finish. However, some major coding environments, such as MATLAB, do not currently support multithreading for customized routines. This limitation in multithreading together with the high system demands of such environments can limit real-time experimental control capabilities. Furthermore, since all tasks are implemented on a single CPU, unresolvable system conflicts may arise if different hardware components are only compatible with certain operating systems.

Network-based parallel processing provides a versatile solution to this problem by dividing experimental tasks across multiple CPUs (***Figure 1C***). In particular, this allows tasks to be executed as parallel processes even if multithreading is not supported. Thus, one major benefit of the REC-GUI framework is that challenges arising from the lack of multithreading support in some high-level programming environments such as MATLAB can be resolved without sacrificing the development benefits of widely used software packages such as Psychtoolbox. This is particularly valuable if computationally demanding real-time stimulus rendering is required. A further benefit not possible with multithreading on a single CPU is that different task components can be implemented using different coding languages and on different operating systems. This feature is especially beneficial since cutting-edge research often requires multiple distinct system components. In the implementation of the REC-GUI framework described here, we highlight this versatility by using MATLAB to render and present stimuli with Psychtoolbox 3 on one CPU, and Python to run a GUI that implements experimental control and behavioral monitoring on a second CPU. With this setup, information about changes in fixation status and user-provided inputs to the GUI are relayed via a network packet to MATLAB which updates the stimulus accordingly. Importantly, this ensures that effectively no cycles of eye position data are lost since sending a network packet takes microseconds. More broadly, network-based parallel processing allows the REC-GUI framework to support a broad range of experimental preparations, as long as the system components support network interfacing.

### Overview of the REC-GUI framework

Experimental control is implemented in the REC-GUI framework using network-based parallel processing. In the implementation described here, experimental tasks are divided into two major groups: (*i*) experimental control and monitoring, and (*ii*) stimulus rendering and presentation (***Figure 2***). However, the number of components and how they are divided can be flexibly determined based on experimental needs. Different groups are executed on separate CPUs that communicate through internet protocols: user datagram protocol (UDP) and transmission control protocol (TCP). The choice of where to use UDP or TCP depends on the task demands. UDP is fast because it does not perform error-checking (processing continues without waiting for a return signal confirming if a data packet was received). TCP is slower than UDP, but extremely reliable for communication because it uses error-checking and temporally ordered data transmission.

**Figure 2.**
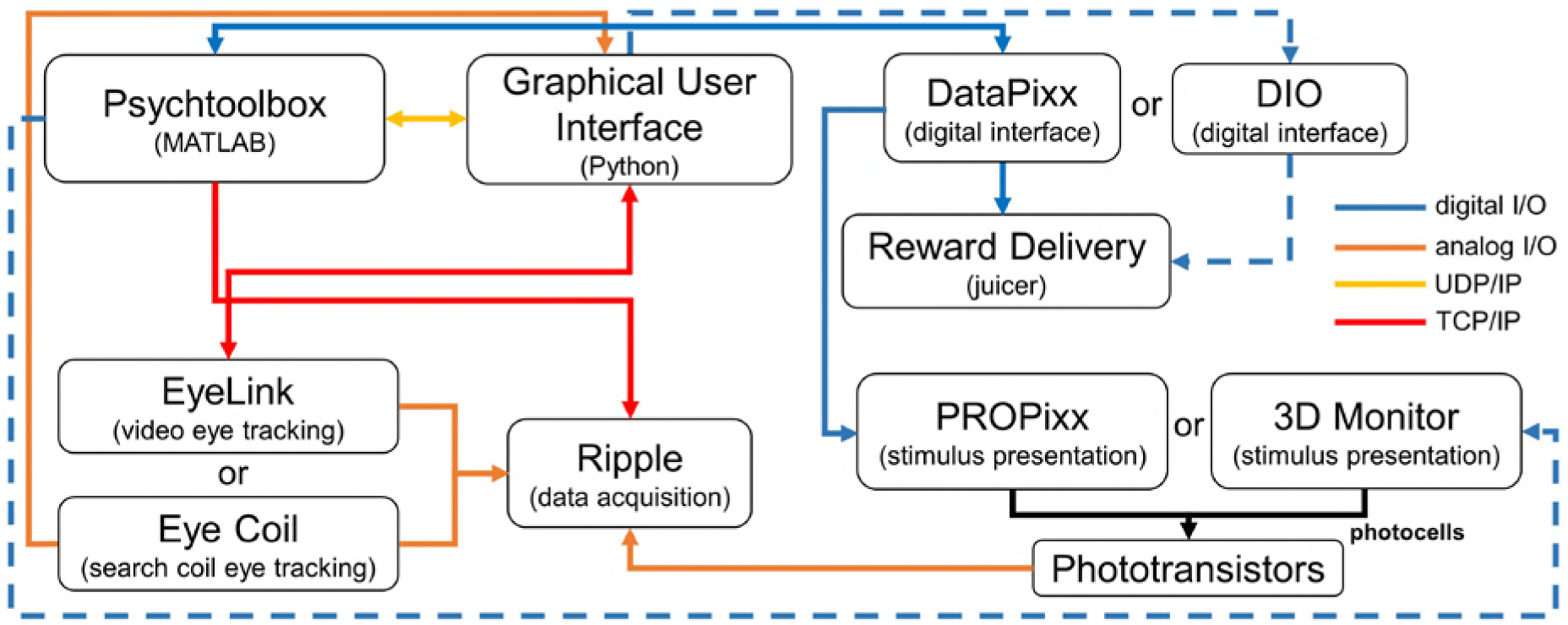
Schematic of the Real-Time Experimental Control with Graphical User Interface (REC-GUI) framework. Experimental control and monitoring is achieved using a GUI that coordinates stimulus rendering/presentation, external devices for measuring behavior and delivering rewards, and data acquisition. Dashed lines show communication pathways for
using non-VPixx displays. Arrows specify the direction of information flow.

For testing, we implemented a system to investigate the neural basis of three-dimensional (3D) visual perception in non-human primates (Rosenberg et al., 2013; Rosenberg and Angelaki, 2014a, b). The experiments require the stereoscopic display of stimuli rendered using multi-view geometry (Hartley and Zisserman, 2003), eye tracking for gaze-contingent stimulus presentation and detecting perceptual reports, reward delivery, and neuronal recordings. Experimental control and real-time monitoring are performed with a GUI coded in Python 2.7 and run on Linux (Ubuntu 14.04, Intel i3 Processor, 8 GB RAM, Intel HD500). Stimulus rendering is performed using Psychtoolbox 3 on MATLAB 2017a, and run on Linux (Ubuntu 16.04, Intel Xeon Processor, 24 GB RAM, NVIDIA GeForce GTX 970). Data acquisition is performed using a Scout Processor (Ripple, Inc.) run on Windows 10 (Intel Xeon Processor, 8 GB RAM, NVIDIA GeForce GT 720).

A detailed schematic of communication between the experimental control and stimulus CPUs during the behavioral task (described below) used for testing is shown in ***Figure 2#x2013;figure supplement 1***. The experimental control CPU uses multithreading to run two major threads for: (*i*) monitoring and evaluating eye position, and (*ii*) updating experimental parameters in real-time based on inputs from the GUI. The stimulus CPU uses a MATLAB while-loop to render stimuli with parameters provided by the GUI, and to achieve gaze-contingent stimulus presentation by repeatedly querying the GUI through a UDP connection established in asynchronous mode so that the while-loop (execution flow) does not pause while waiting for data packets.

### Hardware components

Visual stimuli were rear-projected onto a screen at 240 Hz (120 Hz per eye) using a PROPixx projector (VPixx Technologies, Inc.) and a circular polarizer for stereoscopic presentation. This setup was used for all reported testing because it attains current state-of-the-art limits in 3D display capabilities. A major challenge that arises here is the real-time rendering and high frame rate presentation of 3D stimuli with large depth variations and occlusion without dropped frames or time lags, while enforcing gaze contingencies. Two other setups were used to confirm that the framework is robust to system changes. The second replaced the PROPixx with a VIEWPixx/3D display (VPixx Technologies, Inc.) operating at 120 Hz with active shutter glasses for stereoscopic presentation. The third used a 3D monitor (LG Electronics Inc.) and NVIDIA-2 3D Vision Kit operating at 120 Hz with active shutter glasses (run on Windows 10, Intel Xeon processor, 8 GB RAM, NVIDIA Quadro K4000 graphics card). Results from the two latter setups are not presented here because they confirm the more stringent testing results from the first setup.

The REC-GUI framework supports eye tracking using video or scleral search coil (Judge et al., 1980) methods (***Figure 2***). For the current study, we used video tracking with an EyeLink 1000 plus (SR-Research, Inc.). Binocular eye positions were sampled and digitized by EyeLink, and the measurements sent to the GUI through a TCP connection for real-time analysis. The same measurements were converted into an analog signal and transferred to the Scout Processor on its analog input channels for offline analysis. For the GUI to perform real-time analysis of eye movements measured with search coils, the analog outputs of the coil system would be sampled and digitized using an analog to digital converter (USB-1608G, Measurement Computing, Inc.). The same outputs would be transferred to the Scout Processor for offline analysis.

A Scout Processor (Ripple, Inc.) was used for the acquisition of electrophysiological data and storage of analog and digital signals from external devices. Other acquisition systems that support network interfacing can be substituted with minimal effort. Because timestamps are generated in the Scout Processor, no additional synchronization is required to temporally align input signals. As such, the Scout Processor serves as the main data storage unit, generating and storing a data file with synchronized behavioral and neural signals. The GUI also saves a file containing all data that it transmits and receives along with event codes signaling the occurrence of specific experimental events (e.g., fixation point on, stimulus on) and behavioral events (e.g., fixation acquired, choice made) on the experimental control CPU. Thus, the experimental control CPU provides a backup copy of certain data, and can serve as the main data server for studies that do not have large data demands requiring a standalone acquisition machine.

### System communications

System components communicate through four types of connections: UDP, TCP, analog signals, and transistor-to-transistor logic (TTL) using a digital input/output (DIO). UDP and TCP connections are achieved with network switches (***Figure 2***,**Figure 2–figure supplement 2**), and used for communications between the experimental control CPU, stimulus CPU, EyeLink, and Scout Processor. In this implementation of the REC-GUI framework, two groups of digital event codes are carried over UDP and TCP connections: (*i*) stimulus-related (e.g., fixation target on/off, stimulus on/off, choice target on/off), and (*ii*) behavior-related (e.g., fixation acquired, fixation broken, saccade-to-choice target made). Stimulus-related event codes are triggered in MATLAB and sent to the GUI using UDP and the Scout Processor using TCP (***Figure 2***, ***Figure 2―figure supplement 1***). Currently, there is no Python API for real-time communication between the GUI and Scout Processor. As such, all behavior-related events triggered by the experimental control CPU are sent to MATLAB, which relays event codes to the Scout Processor.

Digital communications are used for a control signal to deliver liquid rewards by sending a TTL pulse to a digital relay output board that opens a solenoid valve. The TTL pulse is either controlled by MATLAB and generated by a DataPixx (VPixx Technologies, Inc.) or controlled by the GUI and generated by a USB DIO interface (USB-1608G, Measurement Computing Inc.), depending on the setup (***Figure 2***).

Experimental systems often require specialized hardware purchased from multiple companies. Network-based communications with that hardware must often occur over non-configurable, predefined subgroups of IP addresses. A simple way to set up communication with such hardware is using multiple parallel networks such that each hardware piece has a single dedicated network switch. In this setup, both the EyeLink and Scout Processor have non-configurable, predefined subgroups of IP addresses that cannot be routed over the same network interface card (NIC). Consequently, two network switches and two NICs are required for both the stimulus and experimental control CPUs, with each NIC assigned to a different subgroup of IP addresses (***Figure 2―figure supplement 2***). With this configuration, both the stimulus and experimental control CPUs can directly communicate with the EyeLink and Scout Processor.

### Configurable experimental control GUI

Information about the ongoing status of an experiment and graphical feedback of critical data is often required to monitor and adjust parameters in real-time using a GUI. To accommodate different experimental paradigms, it is critical that a GUI provides a general method for easily adding/removing parameters from the control set and for manipulating parameters in real-time. To achieve these goals, the REC-GUI framework provides a highly flexible and intuitive user interface for real-time experimental control (***Figure 3***). The GUI displays continuously sampled measurements (e.g., eye position, animal location in an arena, arm position, membrane potential, firing rate, etc.) in the monitoring window. To visually evaluate contingencies, the monitoring window can also display boundary conditions which are used in determining if the measurements fall within a certain criterion range (e.g., if an animal’s gaze is within a certain distance of a fixation target, or if an animal has entered a specific area of an arena). This functionality can also be used to trigger event signals (e.g., TCP or UDP packet, or TTL pulse) to control external devices such as a solenoid, pellet dropper, or neural stimulator. An experimenter can turn boundaries on/off, or change their size, number, locations, etc. in real-time through inputs in the GUI.

**Figure 3.**
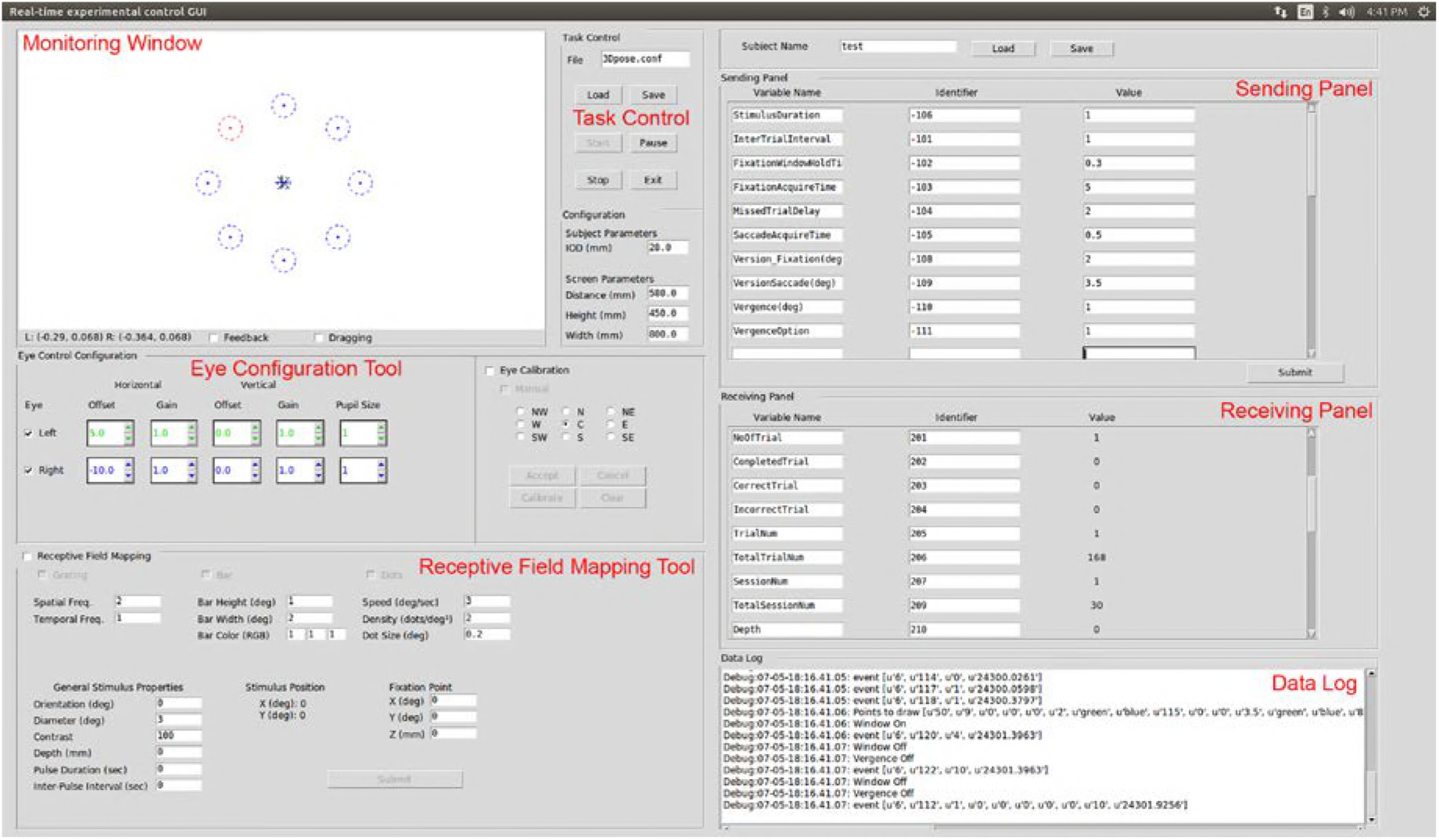
Graphical user interface provided with REC-GUI. The GUI has multiple control panels that are fully customizable. The upper left corner is a monitoring window, here showing a scaled depiction of the visual display. Fixation windows and eye position markers are seen at the center of the monitoring window. Eight choice windows for the 3D orientation discrimination task are also shown (correct choice in red). Below the monitoring window are eye configuration and receptive field mapping tools, which can be substituted for other experiment-specific tools. Task control for starting, pausing, or stopping a protocol is at the center top, along with subject-specific and system-specific configuration parameters. The sending panel in the upper right allows the experimenter to modify task parameters in real time. The receiving panel below that is used to display information about the current stimulus and experiment progress. The lower right panel shows the data log.

Such changes are implemented in the REC-GUI framework using UDP communication to send/receive strings (***Figure 2***, ***Figure 2―figure supplement 1***). Each string in the UDP packet consists an identifier (e.g., −106; a number which the MATLAB code uniquely associates with a specific variable such as ‘stimulus duration’) and a value (e.g., 1 to specify a 1 s duration), followed by a terminator (/qqqq….q padded to 1,024 characters). In this example, the string would be −106 1 /qqqq….q (see User Manual for details). The GUI contains separate panels for sending and receiving UDP packets between the stimulus and experimental control CPUs (***Figure 3***). From the sending panel, an experimenter enters values (e.g., 1) for predefined variables (e.g., stimulus duration) with associated identifiers (e.g., ‘-106’) and clicks ‘Submit’ to send UDP packets to the stimulus CPU. In the current configuration, the sending panel sends control information to the stimulus CPU, but it can also send information to any other machine capable of receiving UDP packets, as required for an experiment. The receiving panel allows the experimenter to predefine identifiers/variables for receiving and displaying information from the stimulus or other CPU (***Figure 3***). In this way, ongoing experimental information can be monitored in real-time. The GUI also contains placeholders in the lower left corner for specialized tools. The default GUI contains interfaces to control eye calibration and receptive field mapping, but these can be easily substituted with other tools. This text-based approach provides a simple way to reconfigure the GUI to meet the demands of different experimental paradigms.

### Animal preparation, behavioral task, and neural recording

The functionality and performance of the REC-GUI framework was tested using a 3D visual orientation discrimination task performed by a rhesus monkey (*Macaca mulatta*). All surgeries and procedures were approved by the Institutional Animal Care and Use Committee at the University of Wisconsin–Madison, and in accordance with NIH guidelines. A male rhesus monkey was surgically implanted with a lightweight Delrin ring for head restraint. At the time of the procedure, the animal was approximately 4.25 years of age and 7.2 kg in weight. After recovery, the animal was trained to fixate a visual target within 2° version and 1° vergence windows using standard operant conditioning techniques.

The animal was trained to perform an eight-alternative 3D orientation discrimination task (***Figure 4***). In the task, the animal viewed 3D oriented planar surfaces, and reported the direction of planar tilt with a saccadic eye movement to an appropriate choice target. Planar surfaces were presented at tilts ranging from 0° to 315° in 45° steps, and slants ranging from 15° to 60° in 15° steps. A frontoparallel plane (tilt undefined, slant = 0°) was also presented. The surfaces were defined as random dot stereograms with perspective and stereoscopic cues (N = 250 dots, each dot subtending ~0.3° of visual angle at the screen distance of 57 cm). Stimuli were centered on the fixation target. At the start of each trial, the animal fixated a target on an otherwise blank screen for 300 ms. The stimulus was then presented for 1,000 ms while fixation was maintained. The stimulus and fixation target then disappeared, and eight choice targets appeared at a radial distance of 11.5° with angular locations of 0° to 315° in 45° increments (corresponding to the possible planar tilts). The animal was rewarded with a drop of water or juice for choosing the target in the direction that the plane was closest to the animal. If fixation was broken before the appearance of the choice targets, the trial was aborted.

**Figure 4.**
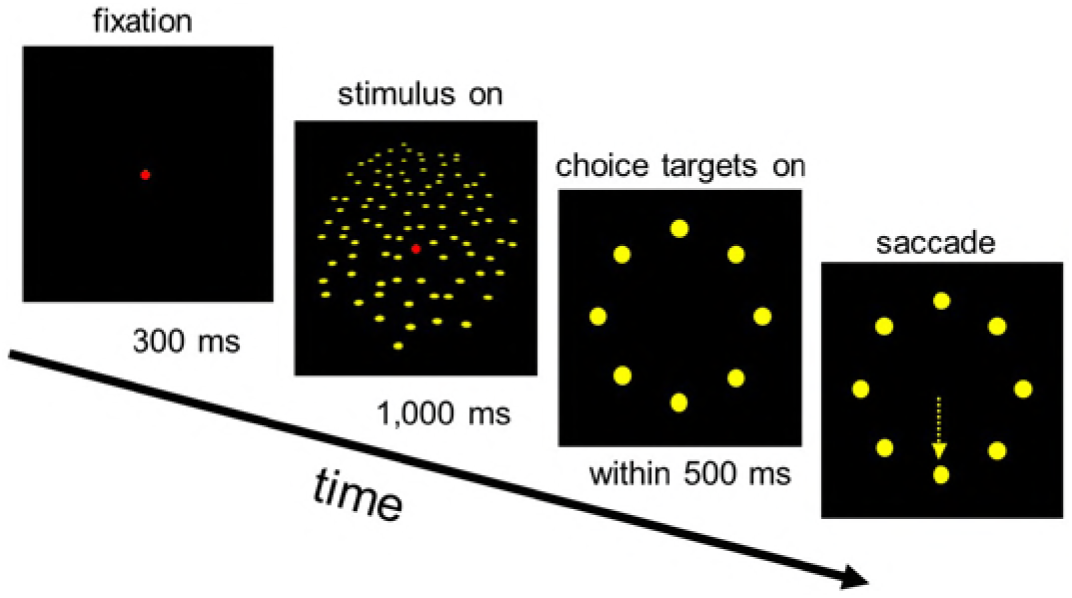
Behavioral task. The animal fixated a target (red) at the screen center for 300 ms. A planar surface was then presented for 1,000 ms (one eye’s view is shown) while fixation was maintained. The plane and fixation target then disappeared, and eight choice targets corresponding to the possible planar tilts appeared. A liquid reward was provided for a saccade (yellow arrow) to the target in the direction that the plane was closest to the animal. Dot sizes are exaggerated and number reduced for clarity.

We measured the 3D orientation tuning of neurons in the caudal intraparietal (CIP) area (Rosenberg et al., 2013; Rosenberg and Angelaki, 2014a, b) while the animal performed this behavioral task. This data was used to confirm the temporal alignment of stimulus presentation, behavioral performance, and neural data in the REC-GUI framework. A tungsten microelectrode (~1MΩ; FHC, Inc.) was targeted to CIP using magnetic resonance imaging scans. The CARET software was used to segment visual areas, and CIP was identified as the lateral occipitoparietal zone (Van Essen et al., 2001; Rosenberg et al., 2013). A recording grid for guiding electrode penetrations was aligned with the brain scan in stereotaxic coordinates using ear bar and grid markers (Laurens et al., 2016). Neuronal responses were sampled and digitized at 30 Khz using the Scout Processor. Single-neuron action potentials were identified by waveform (voltage-time profile) using Offline Sorter (Plexon, Inc.). All subsequent analyses were performed in MATLAB.

## RESULTS

### Network and system performance tests

The REC-GUI framework uses network-based parallel processing to implement experimental control and synchronize multiple experimental devices. In this section, we test the latency of communication between the stimulus and experimental control CPUs resulting from hardware and software processing, as well as the performance of the main loop responsible for stimulus rendering and presentation. Except where otherwise noted, testing was performed with the 3D orientation discrimination task described in the Materials and methods.

Since different experimental processes are implemented on independent CPUs that communicate over a network, it is possible that limitations in network capacity can introduce delays that adversely affect system performance. To test this possibility, we measured the delay resulting from the network hardware. Network latency was measured using a simple ‘ping’ command that is used to test reachability, signal fidelity, and latency in network communication between two hosts (Abdou et al., 2017). This ping sends an internet control message protocol requesting an echo reply from the target host. The latency of the ping is the round-trip duration of the packets between the two hosts. Network latency between the experimental control and stimulus CPUs was found to be negligibly small (average latency = 24 µs; standard deviation: σ = 5.1 µs; N = 1,000 pings), indicating that the network hardware did not introduce substantial delays that could adversely affect performance (***Figure 5A***).

**Figure 5.**
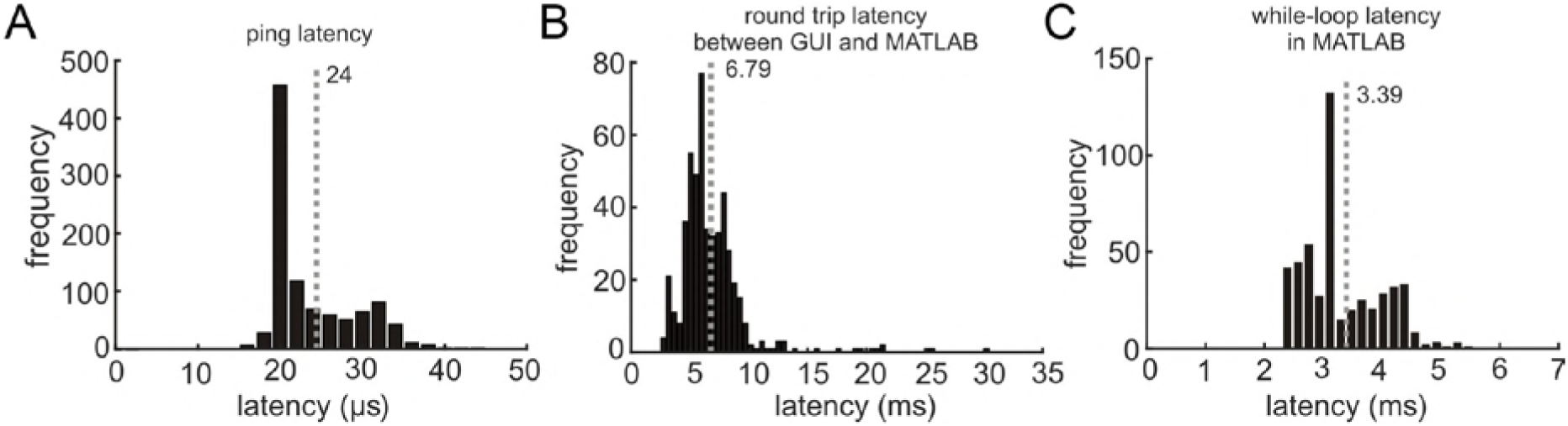
System performance. (A) Ping latency of the network hardware (N =1,000 pings). (B) Overall system performance measured using the round-trip latency of UDP packets between the experimental control (GUI) and stimulus (MATLAB) CPUs during the 3D orientation discrimination task (N =500 round-trip packet pairs). (C) Duration of the main while-loop in the stimulus (MATLAB) CPU for rendering/presenting stimuli (N =3,000 iterations). Vertical gray dotted lines mark mean durations.

Next we measured the overall system performance which includes hardware delays tested above as well as software delays from the main loops (i.e., including all processes executed) on both the CPUs. In this test, the GUI sends a UDP packet to the stimulus CPU and after that packet is detected, MATLAB returns another UDP packet. We measured the latency of multiple round-trip sent/received packets (GUI → MATLAB → GUI), and the distribution of measured latencies is shown in ***Figure 5B***. On average, the total duration of the round-trip packets during the 3D orientation discrimination task was 6.79 ms (σ = 2.9 ms; N = 500). This latency determines the time interval required to synchronize the two CPUs during this computationally intensive task. For many experiments, system performance would likely exceed this level since stimulus preparation in this task was computationally demanding. To determine an upper-bound of system performance with the described REC-GUI implementation, we removed all stimulus-related processes from the MATLAB main while-loop and measured the round-trip latency of the UDP packets. In this case, the round-trip latency dropped to an average of 4.3 ms (σ = 2.3 ms; N = 500). Note that this latency only limits the real-time control of stimulus parameters, and does not limit the precision of the temporal alignment that can be achieved offline, as demonstrated in the neural recording tests described below. Moreover, even with the longer latencies that occurred when rendering complex 3D stimuli, the achieved millisecond-level of control synchronization is sufficient for most experiments.

Lastly, we measured the duration of the main-loop in the stimulus CPU, excluding the UDP processing time for communication with the experimental control CPU. Thus, this test includes all serially executed processes associated with stimulus rendering and presentation, as well as conditional statements for controlling experiment flow. During the 3D orientation discrimination task, the duration of the main-loop was on average 3.39 ms (σ = 1.3 ms; N = 3,000 iterations; ***Figure 5C***), thus accounting for approximately half of the 6.79 ms required to synchronize the two CPUs. To determine a lower-bound duration of the MATLAB main-loop with the same setup, we removed the stimulus-related routines. After removing this code, the latency of the main-loop dropped to an average of 1.7 ms (σ = 0.11 ms; N = 3,000 iterations). Together, these tests show that the performance of the REC-GUI framework can facilitate complex experimental tasks in real-time, with very low latencies using simple, high-level programming environments.

### Stimulus presentation tests

A critical component of experimental studies is the ability to accurately and precisely present stimuli. To evaluate the ability of the REC-GUI framework to present demanding visual stimuli, our tests used large, computationally intensive stereoscopic stimuli presented at 240 Hz (see Materials and methods). The stimuli were rendered as separate right and left eye ‘half-images’ in MATLAB with Psychtoolbox 3, and the ‘flip’ command (Kleiner et al., 2007) was used to alternately present the appropriate image to the right or left eye (120 Hz per eye) for 1 second. To account for the temporal difference between the projector’s refresh rate and the execution rate of the MATLAB script, the flip command was set to wait until the next available cycle of the projector refresh. This setting minimizes the variability in the delay between the flip command and the appearance of the stimulus. To assess the fidelity of the presentation, phototransistor circuits (diagram in the User Manual) were used to track the appearance of stimuli on the screen by detecting a small bright patch in the lower right/left corner of the corresponding (right/left) eye half-images.

The Scout Processor saves the voltage traces generated by the phototransistor circuits to provide a precise signal for aligning events to the stimulus as well as to confirm the fidelity of the stereoscopic presentation on a trial-by-trial basis (e.g., if dropped frames occur on a certain trial, this can be detected and the trial discarded). Example traces showing the presentation of right and left eye images are shown in ***Figure 6A*** (blue and orange traces, respectively). The latency between the initial flip command and the appearance of the stimulus was approximately one cycle of the video refresh (average delay = 4.71 ms, σ = 0.16 ms, N = 500 trials; ***Figure 6B***). The fidelity of alternating right and left eye frames was confirmed by assessing the time lag between the two voltage traces. Since the stereoscopic images were presented at 240 Hz (120 Hz per eye), the right and left eye frame signals should be temporally shifted by ~4.17 ms. We measured the timing difference between each alternation of the right and left eye frames over the 1 s stimulus duration for 500 trials. The histogram of timing differences shows a strong peak at 4.16 ms (minimum = 3.9 ms; maximum = 4.4 ms; σ = 0.066 ms), indicating that the right and left eye frames were well synchronized at the intended 240 Hz stimulus presentation rate (***Figure 6C***). This histogram also confirms that no frames were dropped over the cumulative 500 s of stimulus presentation (dropped frames would appear as timing differences ≥ 12.5 ms).

**Figure 6.**
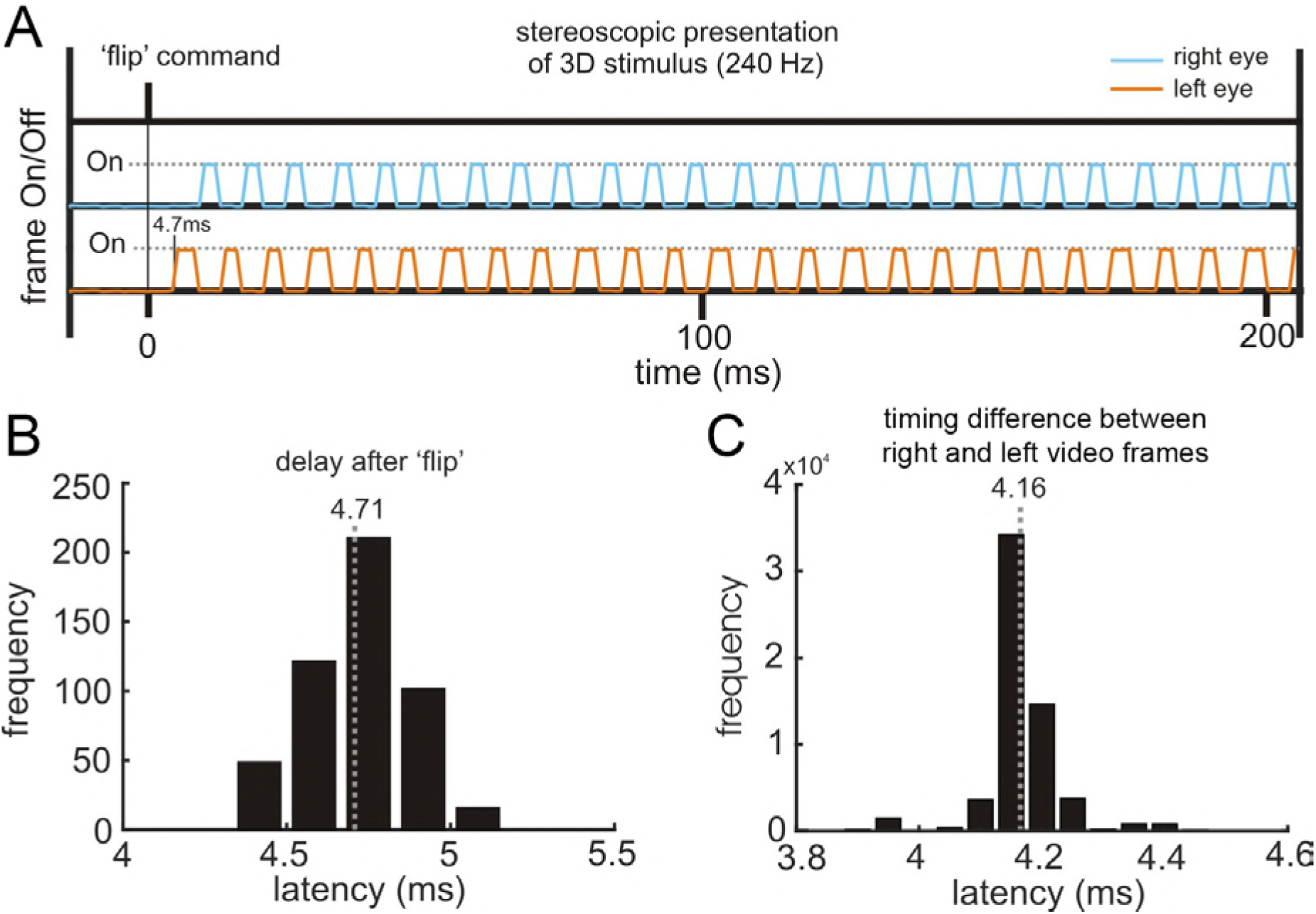
Quantification of the fidelity of stimulus presentation. (A) Right and left eye frame signals measured on the screen. The anti-phase rise and fall of the two signals indicates that the right and left eye signals are temporally synchronized. (B) Latency between the initial flip command in MATLAB and the appearance of the stimulus (N = 500 trials). Approximately one frame occurs between the flip command and stimulus appearance. (C) Histogram of timing differences between the two eyes’ frames peaks at 4.16 ms indicating that 240 Hz (120 Hz per eye) stimulus presentation is reliably achieved (data from 500 trials). Voltage traces were sampled at 30 kHz. Mean times are indicated by vertical gray dotted lines.

### Real-time experimental control

Real-time monitoring to guide the control of ongoing processes is critical for many experimental studies. We evaluated this capability of the REC-GUI framework using gaze-contingent stimulus presentation. Right and left eye positions were sampled at 1kHz using an EyeLink 1000 plus (SR-Research Inc.). The eye position data were fed to the GUI which implements routines for evaluating if the animal is holding fixation on the target, if fixation is broken, or if a particular choice is made in the 3D orientation discrimination task. Depending on the results of these routines, the experimental control CPU then directs the stimulus CPU to enter particular experimental stages (e.g., fixation only, stimulus presentation, choice targets, etc.).

We first confirmed that the GUI successfully enforced version and vergence eye position during a fixation task. Fixation targets were presented at three distances relative to the viewing screen, and the task required that fixation be maintained for 1 s. The top row of ***Figure 7A*** shows right and left eye version enforcement windows along with eye traces for 4 representative successfully completed trials at each distance. The bottom row of ***Figure 7A*** shows the vergence errors for the same trials. Second, we confirmed the successful enforcement of gaze relative to the fixation target during the 3D orientation discrimination task. ***Figure 7B*** shows horizontal and vertical eye displacements for each eye for 34 representative successfully completed trials. Lastly, ***Figure 7C*** shows the full eye traces for the same 34 trials, showing that saccadic eye movements to each of the eight choice targets were accurately detected.

**Figure 7.**
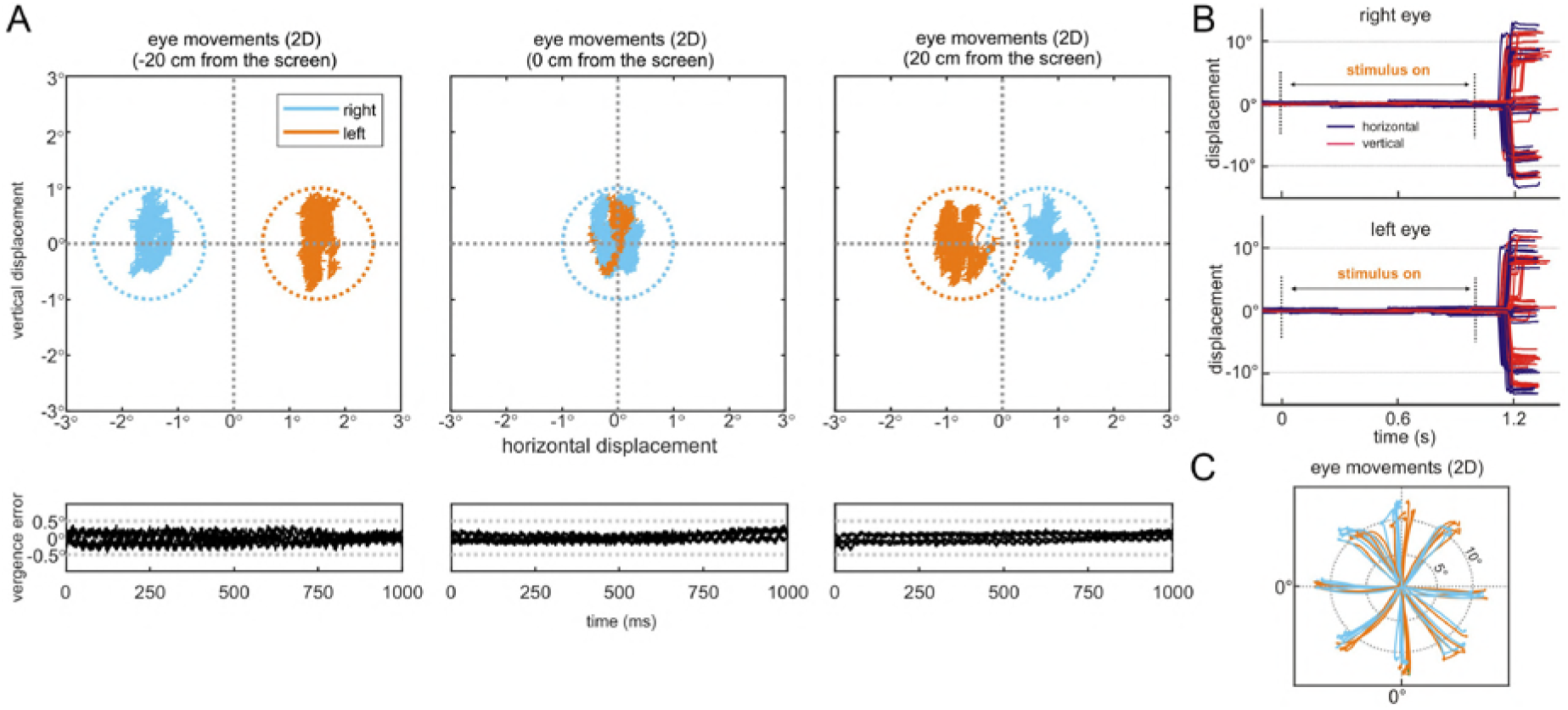
Verifying behavior-contingent stimulus presentation. (A) Enforcement of version and vergence during fixation at different distances. Top row shows left and right eye traces while fixation was held at three different distances using stereoscopically rendered targets (N =4 trials/distance). Circular windows show 2° version enforcement windows. Bottom row shows simultaneously measured vergence error traces. Gray dotted lines show a 1° vergence enforcement window. (B) Right and left eye traces during the 3D orientation discrimination task, aligned to the stimulus onset (N =34 trials). The horizontal (vertical) component of the eye movements are shown in purple (red). (C) Eye movement traces showing the choice target selection (same data as in B).

### Temporal alignment of neural data to stimulus-related and behavioral events

To confirm the ability to precisely align events in time, we measured the 3D surface orientation tuning of a CIP neuron while the animal performed the 3D orientation discrimination task. First, we confirmed the ability to precisely align stimulus-driven neuronal responses to the stimulus onset detected by the phototransistor. A raster plot showing the timing of individual action potentials for 545 trials aligned to the stimulus onset (each row shows a different trial) along with the spike density function (red curve; convolution with a Gaussian function of σ = 10 ms) (MacPherson and Aldridge, 1979) is shown in ***Figure 8A***. The precise alignment of spike times relative to the stimulus onset reveals a sharp visual transient in the spike density function. Second,

**Figure 8.**
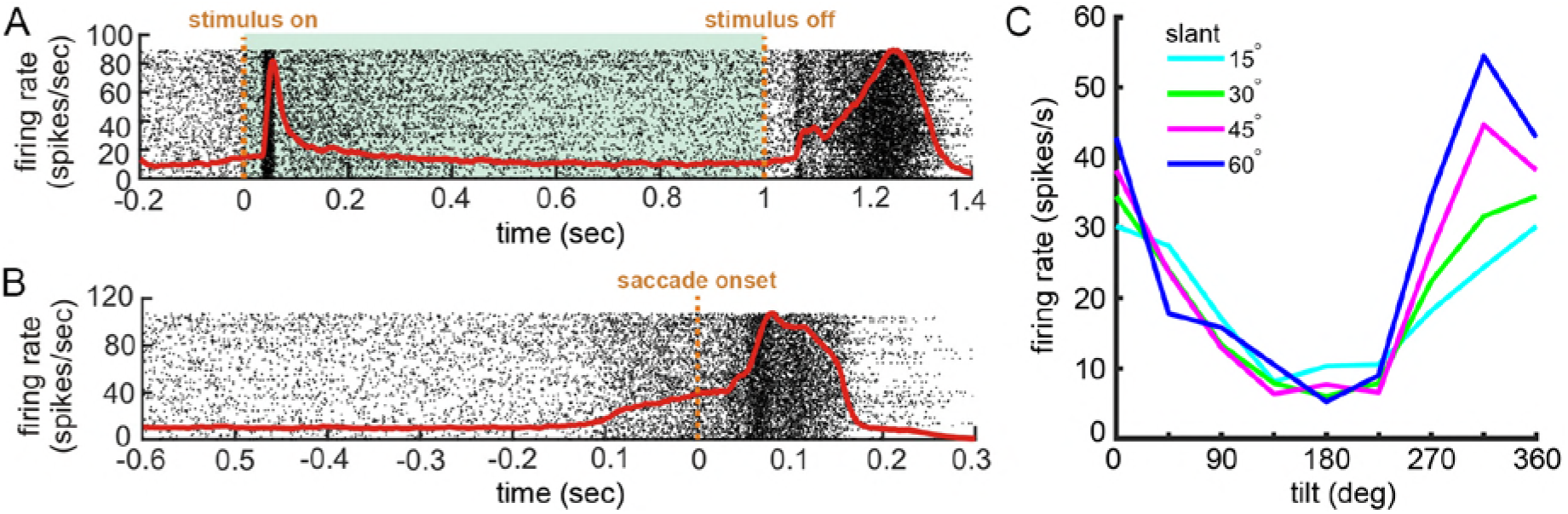
Temporal alignment of neuronal responses to stimulus-related and behavioral events. (A) Raster plot showing the timing of action potentials aligned to the stimulus onset (N = 545 trials). Shaded region marks the stimulus duration. (B) Raster plot showing the timing of action potentials aligned to the saccade onset. Each row is a different trial, and each dot marks a single action potential. Red curves are spike density functions. (C) 3D orientation tuning. Each curve shows tilt tuning at a fixed slant.

We confirmed the ability to precisely align neuronal responses to the measured choice saccades. Saccade onsets were detected offline as the first time point at which the eye movement was faster than 150°/s (Kim and Basso, 2008, 2010). Spike times were aligned to the saccade onset and the spike density function calculated (***Figure 8B***). Note the build-up of neuronal activity preceding the saccade that was not evident when the responses were aligned to the stimulus onset. As expected in CIP (Rosenberg et al., 2013), the neuron was jointly tuned for slant and tilt (***Figure 8C***).

## DISCUSSION

Test results confirm that the REC-GUI framework provides an accurate and precise solution for implementing demanding neuroscience studies with millisecond-level control. By achieving robust experimental control with high-level programming environments, technical challenges that hinder labs from conducting complex, behaviorally relevant research can be overcome without the need for low-level programing languages or professional programmers. Beyond more traditional experiments with a fixed trial structure, the REC-GUI framework capabilities can support research involving the use of naturalistic, complex stimuli that are dynamically updated based on real-time behavioral or neuronal measurements. For example, the system can support closed-loop experiments in which multisensory visual–vestibular stimuli are updated based on active steering behaviors rather than passive, predefined motion profiles. Alternatively, stimuli can be updated or neural activity perturbed through electrical or magnetic stimulation with very short latencies triggered by real-time behavioral or neuronal measurements. Such experiments will be critical to understanding the relationship between the dynamic activity of neural populations, perception, and action during natural behaviors. The REC-GUI framework can facilitate such research by reducing the overhead associated with parlaying the necessary technology into a cohesive experimental system. By reducing technical hurdles and providing a flexible experimental framework, complex experimental protocols will be accessible to a greater number of labs, and the opportunity for scientific discovery increased.

Towards this goal, the REC-GUI framework offers several advantages compared to other control systems. First, network-based parallel processing makes the framework inherently modular and highly flexible. Since the only constraint on incorporating specialized software or hardware is that network interfacing is supported, a broad range of devices for stimulus presentation (visual displays, speakers, pellet droppers, etc.), behavioral measurement (button presses, eye movements, biometrics, location, etc.), and other experimental needs (stimulator, osmotic pump, etc.) can be used out of the box. Such flexibility allows the framework to be adapted to a broad range of experimental preparations (*in vitro*, anesthetized, or awake-behaving) and neuroscience research domains (sensory, cognitive, motor, etc.), and makes it agnostic to the type of neural data recorded (electrophysiological, optical, magnetic resonance imaging, etc.). For instance, we also use the REC-GUI framework to conduct perceptual learning studies with adolescents with autism. Second, dividing computing demands across CPUs improves system performance and enables researchers to implement system components using different coding languages and operating systems. This reduces compatibility issues and increases the efficiency with which experimental setups are configured. Likewise, while the performance of the REC-GUI implementation presented here is sufficient for most experimental needs, if greater precision is required, this can be achieved using a lower-level programming language such as C or C++ for stimulus rendering and presentation. Third, diverse data types can be collected in parallel at different temporal resolutions, and precisely aligned, allowing for multi-faceted research. Lastly, a user-friendly, customizable GUI allows for intuitive and flexible experimental design changes.

A number of other control systems exist, so we briefly review some of them and benchmark the REC-GUI framework with them to help facilitate system selection. The Laboratory of Sensorimotor Research (LSR) real-time software suite from the National Eye Institute provides a full package including a user interface, visual stimulus rendering, data acquisition, and offline data analysis tools (Hays et al., 1982). Similar to REC-GUI, the LSR suite divides experimental control, stimulus processing, and data acquisition across CPUs. The LSR suite is highly powerful, and can meet the demands of many complex behavioral paradigms. However, the LSR scripting language is relatively complex, making the overhead of learning the system and developing new experiments quite high. By using high-level programming environments, the REC-GUI framework aims to reduce this overhead, particularly as it relates to modifying or adding new experimental tasks. Additionally, the LSR suite is specialized for visuomotor studies, whereas the REC-GUI framework is agnostic to the subdomain of neuroscience research.

A freely available system that is fully implemented in MATLAB is MonkeyLogic (Asaad et al., 2013). The system achieves millisecond-level temporal resolution, and provides a user interface with real-time behavioral monitoring. In addition, it is convenient to implement control flows for new behavioral paradigms. MonkeyLogic is designed for a single CPU, so experimental control is performed serially due to MATLAB’s multithreading limitations. This can be problematic for real-time control when the stimulus rendering/presentation demands are high. For example, since stimuli are transferred to the video buffer during the inter-trial interval without compression, it may not be suitable for presenting long-duration stimuli at high frame rates. For instance, using MonkeyLogic for the experiment implemented here would result in long inter-trial intervals to transfer the stimuli to the video card, dropped frames, and presentation lags. Along this line, the MonkeyLogic forum indicates that it cannot support 240 Hz visual stimulus presentation (http://forums.monkeylogic.org/post/high-refresh-rates-vpixx-8408242). Similar limitations will exist for other single CPU, MATLAB-based control systems (e.g., PLDAPS), though some limitations may be at least partially remediated if the real-time monitoring and control features provided by a GUI are eliminated (Eastman and Huk, 2012). Such systems may be ideal for tasks that do not have high real-time behavioral contingency and stimulus rendering/presentation demands, since network communications make the REC-GUI framework slightly more complex. Additionally, there is some added cost to setting up the REC-GUI framework compared to single CPU systems since it requires multiple CPUs, but that cost difference is relatively small. In exchange, the REC-GUI framework allows high-level programming environments to be used to robustly control computationally demanding and behaviorally complex neuroscience experiments.

To facilitate customization and future developments, we provide sample MATLAB scripts and Python GUI code (https://rosenberg.neuro.wisc.edu/). We will maintain the REC-GUI framework as an open-source project, and welcome development contributions from others. Our hope is that this will help researchers perform multi-faceted research combining physiology and behavior by reducing time spent solving technical problems, and increasing time focused on experimental questions and design. The REC-GUI framework will also help promote research transparency, standardize data acquisition, and improve reproducibility by facilitating cheap and easy replication of experimental paradigms.

---

**Figure 2 – figure supplement 1.**
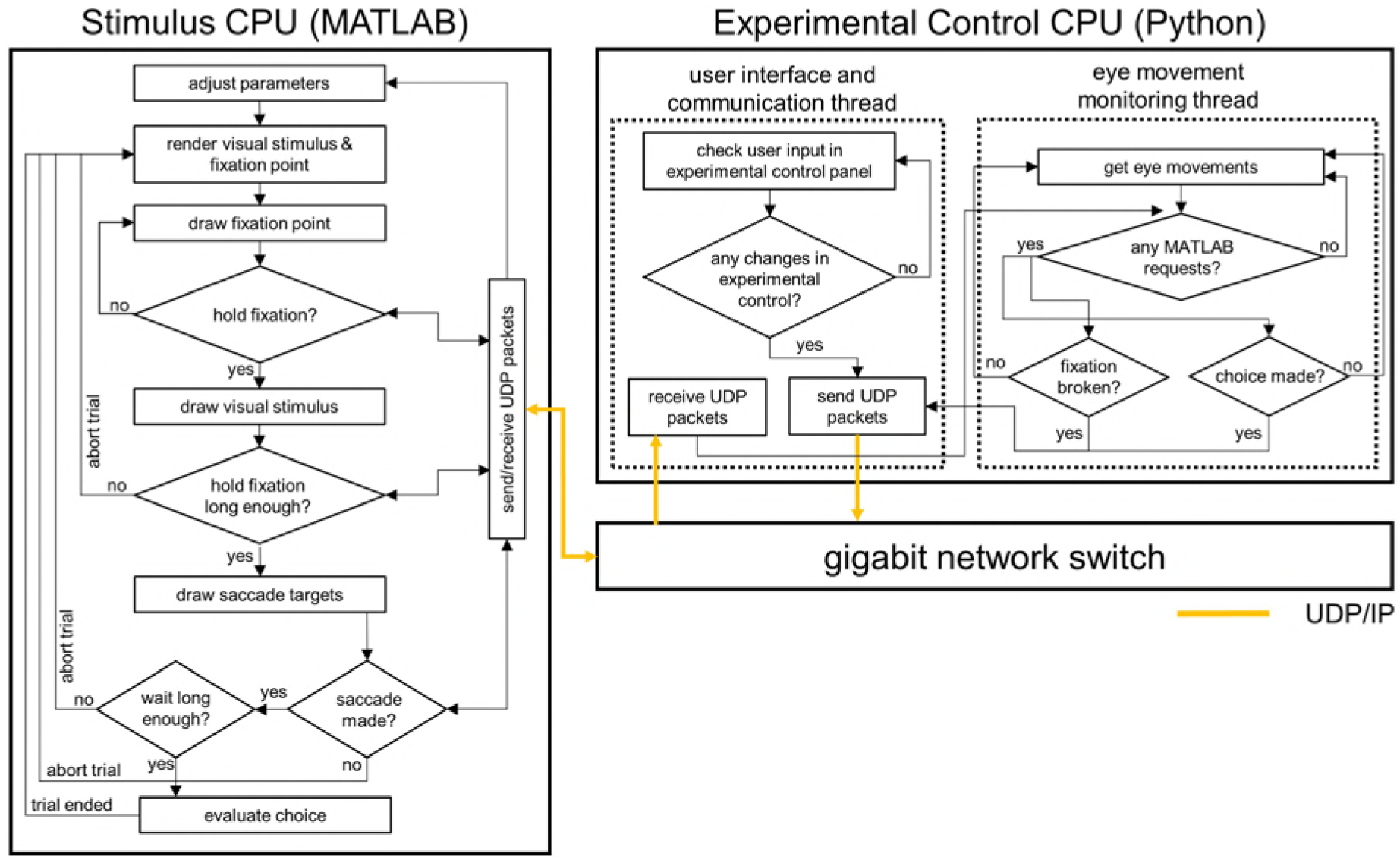
Communication flowchart between the stimulus CPU (MATLAB) and experimental control CPU (Python) showing the exchange of experimental parameters for stimulus rendering and behavioral control. Arrows specify the direction of information flow.

**Figure 2 – figure supplement 2.**
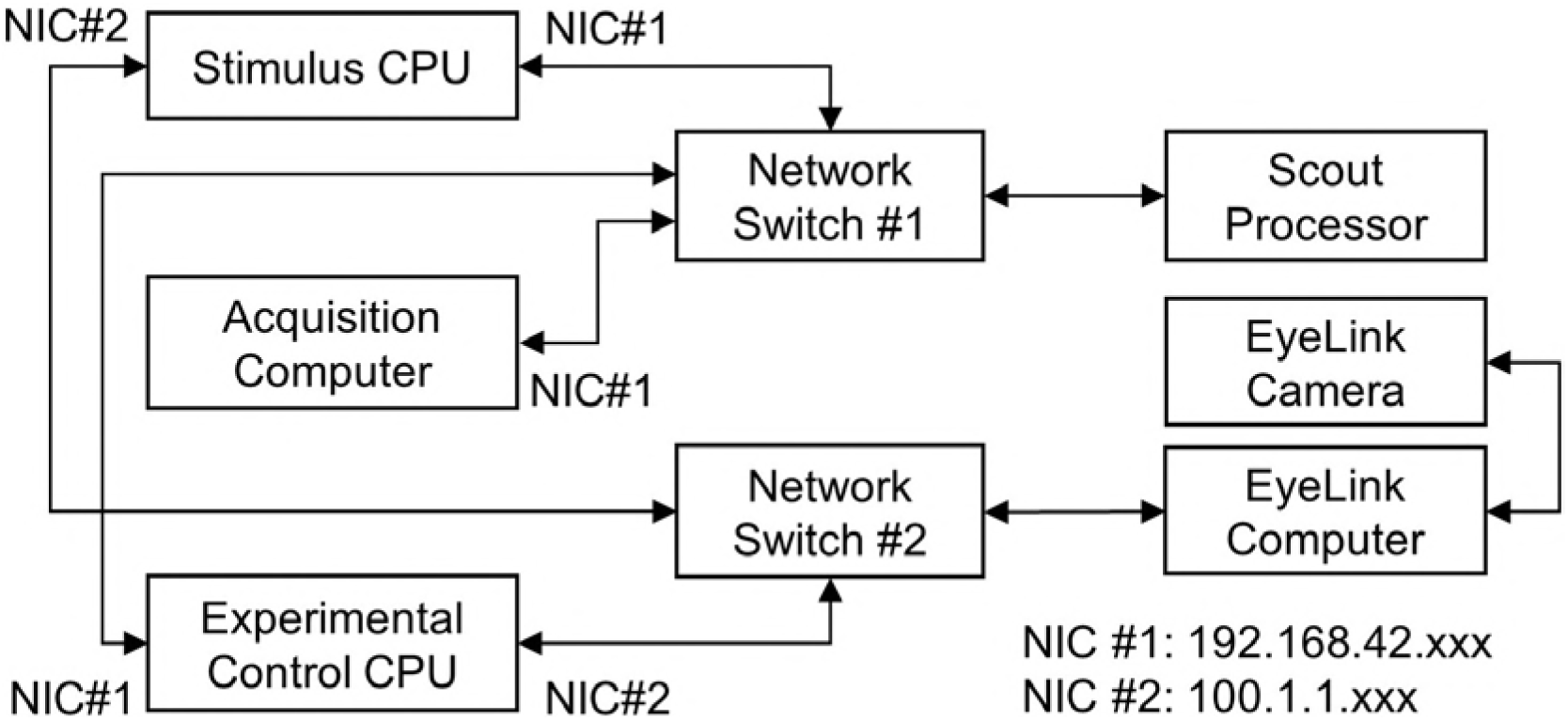
Network configuration. The Scout Processor and EyeLink have non-configurable IP addresses, so two network switches are required to route two different subgroups of IP addresses to the stimulus and experimental control CPUs. Arrows specify the direction of information flow.

## Acknowledgments

This work was supported by the Alfred P. Sloan Foundation, Whitehall Foundation Research Grant 2016-08-18, and National Institutes of Health Grants DC014305 and EY029438. Further support was provided by National Institutes of Health Grant P51OD011106 to the Wisconsin National Primate Research Center, University of Wisconsin – Madison.

## Notes

**Competing Interests:** The authors have no competing financial or non-financial interests.

